# The defects of the hippocampal ripples and theta rhythm in depression, and the effects of physical exercise on their amelioration

**DOI:** 10.1101/2023.01.30.526359

**Authors:** Shinnosuke Koketsu, Kohki Matsubara, Yoshino Ueki, Yoshiaki Shinohara, Koichi Inoue, Satona Murakami, Takatoshi Ueki

**Affiliations:** Department of Rehabilitation Medicine, Nagoya City University Graduate School of Medical Sciences, Nagoya, Aichi, 467-8601, Japan; Department of Integrative Anatomy, Nagoya City University Graduate School of Medical Sciences, Nagoya, Aichi, 467-8601, Japan; Department of Physical therapy, Nagoya Women’s University Faculty of Medical Science, Nagoya, Aichi, 467-8601, Japan; Department of Anatomy and Cell Biology, Yamanashi University Graduate School of Medical Sciences, Chuo, Yamanashi, Japan

## Abstract

Accumulated evidence demonstrate that environmental stress affects the hippocampus, functioning in cognition and sociality, and causes various depressive symptoms. In addition, recent findings showed that environmental stress influenced the hippocampal activity correlated with neuroinflammation, and impaired the hippocampal sharp wave ripples (SWRs), pattens of spike sequences, and the theta rhythms, a strong oscillation observed in the hippocampus. The involvement of the electrophysiological alterations in the etiology of depression has not been appreciated especially in the hippocampus. Furthermore, the pathological markers associated with such alterations have not been identified. In the present study, therefore, the impairment of the SWRs and the theta rhythms in the hippocampus of the restraint stress-induced depression model of mice was analyzed. In the model mice the hippocampal SWRs and theta rhythms were impaired in depression, while physical exercise significantly reverted them. As previously reported, chronic stress induced inflammation in the affected hippocampus in parallel with defects of adult neurogenesis, on the other hand physical exercise ameliorated those pathological conditions of the bran in depression. In conclusion, this study demonstrated the implications of impairment of the hippocampal SWRs and theta rhythms in the etiology of depression and their usefulness as diagnostic markers of depression.

## 1. Introduction

The pathophysiology of depression remains to be elucidated, although a number of associated vulnerable genes and environmental cues have been identified[1,2]. On the other hand, several kinds of therapeutic interventions for depression have recently been demonstrated to ameliorate its psychiatric symptoms and improve the patient’s quality of life. Among those, physical exercise mostly has been proved to alleviate characteristic symptoms of depression such as social disorder and cognitive defect. Therapeutic targets of physical exercise in the affected brain in depression have not been precisely identified, however, recent studies revealed that hippocampal neural connections with the prefrontal cortex, amygdala, and hypothalamus, which function in sociality and anxiety, were impaired[3,4], and furthermore exercise strengthens hippocampal function, associated with sociality and cognition[5]. In addition, accumulated evidence suggested that inflammation in those brain areas was involved in the etiology of hippocampal dysfunction in depression[6,7]. These findings lead to the idea that electrophysiological examination such like electroencephalogram of the alteration of hippocampal activity in depression is applicable for the diagnosis of neurological impairment related to the pathophysiology of depression. The authors recorded sharp wave ripples (SWRs) and theta rhythm in the hippocampus in mouse model of depression and examined whether SWRs and theta rhythm reflect the neurological states characteristic of depression, and whether exercise influences SWRs and theta rhythm as the result of its improvement of depressive symptoms.

SWRs, patterns of spike sequences thought to be involved in memory consolidation, are local field potentials, which reflect transient excitatory drives and high-frequency oscillations, based on pyramidal cell-interneuron interactions in the hippocampus[8,9]. SWRs generate while resting and sleeping, which implies that environmental stress reduces the generation of SWRs[10]. Machineries generating SWRs and their regulatory neural bases have not been understood in detail, and their correlation with the pathophysiology of neurological and psychiatric disorders have not been appreciated to date. On the other hand, the hippocampal theta rhythm occurs on the basis of projection from the medial septum, which receives inputs from the hypothalamus and the brain stem. The hippocampal theta rhythm has recently been reported to occur during REM sleep and link with arousal, while other studies demonstrated that theta rhythm functions in navigation, learning and memory. The correlation of hippocampal theta rhythm with the causative input from the hypothalamus, which plays a crucial role in motivation, evokes its involvement in the etiology of depression.

In the present study, the influence of physical exercise on behavioral symptoms of depression, and the correlation of the alteration of SWRs and theta rhythm with the amelioration of depressive symptoms were examined in the mouse model of depression. As a result, exercise improved depressive behaviors, while SWRs and theta rhythm restored in parallel with the alleviation of depression. Of note is that the alteration of SWRs and theta rhythm cooccurs, accompanying the hippocampal neuroinflammation, which underlies the etiology of the cognitive impairment. In conclusion, our findings demonstrate that alteration of hippocampal SWRs and theta rhythm links with the amelioration of depressive symptoms, where neuroinflammation simultaneously reduces, and hippocampal SWRs and theta rhythm are useful as diagnostic indexes to estimate the severity of symptoms in depression.

## 2. Methods

### 2.1 Animals

45 healthy male C57BL/6 mice (8 week-old) were purchased from Japan SLC. Mice were housed in a group of up to five animals on a 12h light/dark cycle at 22◻ with free access to food and water. The handling, use of animals, and experimental protocols were approved by the Institutional Animal Care and Use Committee of Nagoya City University. All experimental procedures were conducted according to ARRIVE guidelines. All methods were performed in accordance with the guidelines from American Veterinary Medical Association and our institution.

### 2.2 Chronic restraint stress and spontaneous exercise

Mice were randomly divided into 3 groups, 1) a group of healthy control mice (HC, n=15), 2) a group of mice subject to chronic restraint stress (CRS, n=15), 3) a group of mice subject to chronic restraint stress and physical exercise (ExCRS, n=15), and were bred in each environment for 21 days. HC mice were housed in a 20 x 25cm plastic cage. CRS mice were subject to chronic restraint stress, which was previously described[11–13]. Briefly, individual mouse was inserted into a 50 ml tube with small holes on its wall, plugged up with pieces of paper towel, and left for 4 hours. For the rest hours, CRS mice were bred similarly with HC mice. On the other hand, ExCRS mice were subject to the same stress and housed in 25 x 30cm plastic cage with running wheel.

### 2.3 Forced swim test

The test was performed according to previous studies[11,13–19]. The mice were forced to swim for 6 min in a cylindrical vessel with 30 cm height and 10 cm diameter, filled with water at 24 ◻. In the test, the movement of the mice was captured with video camera, and the total hours of immobility, in which the mice floated with minimal movement, were quantitatively analyzed.

### 2.4 Open field test

The test was conducted in a white closed box (40 x 40 x 40 cm). The test was performed according to previous studies[11,12,16,17,20,21]. Mice were gently placed in the center of the box and freely explored. All the sessions were recorded on video and captured on a PC for analysis. A 20 cm square in the center of the box floor was defined as the center region. The time period the animal spent in the center region, number of times it entered the center region, and the distance it moved were analyzed. At the end of the session, the box was thoroughly cleaned with ethanol and dried.

### 2.5 Novel object recognition test

The novel object recognition test was performed to examine the hippocampus-dependent cognitive function, based on the tendency of animals to have interests in new objects and spend more time for exploration[17,22–24]. At the beginning, two identical objects were placed in the box, and a mouse was allowed to explore the box for 10 min (learning step). After 2 hours, one of the two objects was replaced with a new color and shape object and a mouse was again allowed to explore it for 10 min (memory step). All the sessions were recorded on video, and the duration exploring the new object was measured as an evaluation index.

### 2.6 Electrophysiology

Local field potential (LFP) recording was performed as previously described[25,26]. Briefly, the mice were anesthetized with urethane (1.2-1.4 g/kg) and fixed in a stereotaxic frame (Narishige). After removing the scalp, a hole was drilled above the dorsal hippocampus on either side (Bregma: mediolateral 2.0 mm, anteroposterior −2.0 mm). The dura was surgically removed, and a 16-channel linear silicon probe (inter-channel distance = 50 μm; Alx15-5 mim-50-177-A16; NeuroNexus) was slowly inserted to the hippocampal CA1 so that the middle channel was located in the stratum pyramidale. A mixture of white petrolatum and liquid paraffin was applied to the surface of the brain to prevent it from drying out. Wideband (0.1-9000 Hz) extracellular field potentials were continuously recorded with a sampling rate of 31 kHz (TDT). Rectal temperature was measured and a heat mat was used to maintain the body temperature at 37 ◻.

### 2.7 LFP data analysis

LFP data analysis was performed, using MATLAB (MathWorks Inc.) as was reported previously[25,26]. To analyze the ripple event, LFP signals in the stratum pyramidale were first resampled to 20 kHz. LFP signals were band-pass filtered over the ripple frequency band (80-250 Hz) and the resulting signals were squared and smoothed with a 19.2 ms length humming window. In the initial screening, the ripple events were detected as a period, during which the smoothed signal exceeded the mean of 7 times SD at a ripple interval of 100 ms. In the second screening, a minimum value within ± 35 ms from each detected point (ripple through) was assigned as the ripple timing, and a 200 ms ripple filtering waveform centered on the ripple timing was extracted and further analyzed. After automatic detection, visually obvious noises were manually removed from further analyses. Ripple duration was defined as the range surrounding the ripple timing, in which the amplitude of the waves continuously more than doubled the SD. The peak amplitude of ripples is defined as the maximum amplitude through the extracted ripple event and expressed in absolute value. The frequency of ripples during non-theta periods was examined. Ripples observed within 35 ms were considered as one ripple event, and the sum of ripple events was considered as unilaterally occurred ripple.

The original data of LFP was resampled to 1.25 kHz and the spectrogram was calculated to automatically detect theta period. The theta period was detected in the LFP recorded from the CA1 radial layer in the right hemisphere (150 μm below the stratum pyramidale). The theta period was detected in accordance with following criteria: 1) The ratio of the peak power of the theta band (3.5 - 7 Hz) to that of the delta band (2 - 3 Hz) exceeds 0.6. 2) The period is more than 10 s. Furthermore, the power spectral density of the detected theta period was estimated by the Welch periodogram method. The theta, low gamma and high gamma powers were calculated by integrating the power spectral densities of 3 - 6, 40 - 55, and 65-90 Hz, respectively. Spectral power was calculated from the average of the data of animal experiments, which was obtained from the multiple recording sessions.

### 2.8 Histology and immunohistochemical analysis

The mice were anesthetized with urethane and transcardially perfused with 4% paraformaldehyde in 0.1 M phosphate buffer (PB). The brain samples were cut out, post-fixed overnight at 4 ◻, submerged in 30% sucrose in 0.1 M PB, and then frozen at −80 ◻. The 10 μm slices of the hippocampus and cerebral cortex were prepared by the cryostat (Thermo Fisher Scientific). Immunohistochemical staining was performed as previously reported[27]. Briefly, the brain sections were permeabilized with 0.1% Triton X-100 in PB for 20 min, blocked with 5% bovine serum albumin for 30 min, and incubated overnight with primary antibodies: anti-Sox2 (rabbit IgG, 1:250, ab97959, Abcam), anti-ΔFosB (rabbit IgG, 1:500, #2251, Cell Signaling Technology), anti-Iba1 (rabbit IgG, 1:500, 013-26471, Wako). As a secondary antibody, anti-rabbit IgG-Alexa Fluor 594 were used. After counterstaining with DAPI (Vector Laboratories), the tissue images were taken under fluorescent microscope (Olympus). The numbers of the cells expressing Sox2, ΔFosB, and Iba1 were quantified by randomly choosing 4 view fields (100 x 100 μm each) under the microscope and taking an average of the numbers of the cells. Morphological analysis of microglia was performed by skeleton analysis method. Briefly, 10 images of microglia were cropped from each image and converted to 8-bit grayscale images, binarized and skeletonized using ImageJ. Skeleton analysis of ImageJ plug-in was performed in the obtained images, and the number and the total lengths of the processes were automatically analyzed[28,29].

### 2.9 Statistics

Unless otherwise noted, comparisons of multiple populations were performed using a one-way ANOVA followed by the Tukey-HSD test (MATLAB 2019b). A p-value 0.05 was considered as significant.

## 3. Results

### 3.1 Development of restraint stress-induced depression model of mouse, and the effect of physical exercise on the depressive symptoms

The mouse model of depression was developed by restraining the mouse in a cramped container for 4 hours every day over a 3-week period (chronic restraint stress (CRS) mice), while the healthy control (HC mice) was reared in normal housing. In addition, the effect of physical exercise on restraint stress-induced depression was examined in the mice, which were subject to the restraint stress for 4 hours every day and subsequent voluntary exercise over a 3-week period (exercise and chronic restraint stress (ExCRS) mice) (Fig. 1A, B). After rearing for 3 weeks, change of body weight and depressive behavior were analyzed in CRS, HC and ExCRS mice. The body weight significantly decreased in CRS mice compared to that in HC mice, and reverted considerably in ExCRS mice (HC-CRS, p<0.001; HC-ExCRS, p<0.001; CRS-ExCRS, p=0.048) (Fig.1C). The forced swimming test to estimate depressive behavior[15,18,19] demonstrated that the total immobility time for the period of 6 min significantly extended in CRS mice compared with that in HC mice, and shortened in ExCRS mice (HC-CRS, p=0.0032; CRS-ExCRS, p=0.0012) (Fig.1D). The open field test showed that the number of times to enter to the center and the time period to spend in the center of the field significantly decreased in CRS mice compared to those in HC mice (entries; HC-CRS, p<0.001; CRS-ExCRS, p<0.001: time of stayed; HC-CRS, p<0.001; CRS-ExCRS, p<0.001) (Fig.1E). The result demonstrated that physical exercise considerably attenuated anxiety in the depression model mice[20,21]. The novel object recognition test was performed to assess the hippocampal cognitive function[22–24], and the result showed that the ratio of the duration exploring novelty object to the total duration of exploration significantly decreased in CRS mice compared with HC mice, while that ratio reverted in ExCRS mice (HC-CRS, p=0.0037; CRS-ExCRS, p=0.042) (Fig.1F). The data suggest that physical exercise ameliorates the hippocampal cognitive dysfunction in depression.

**Figure 1.**
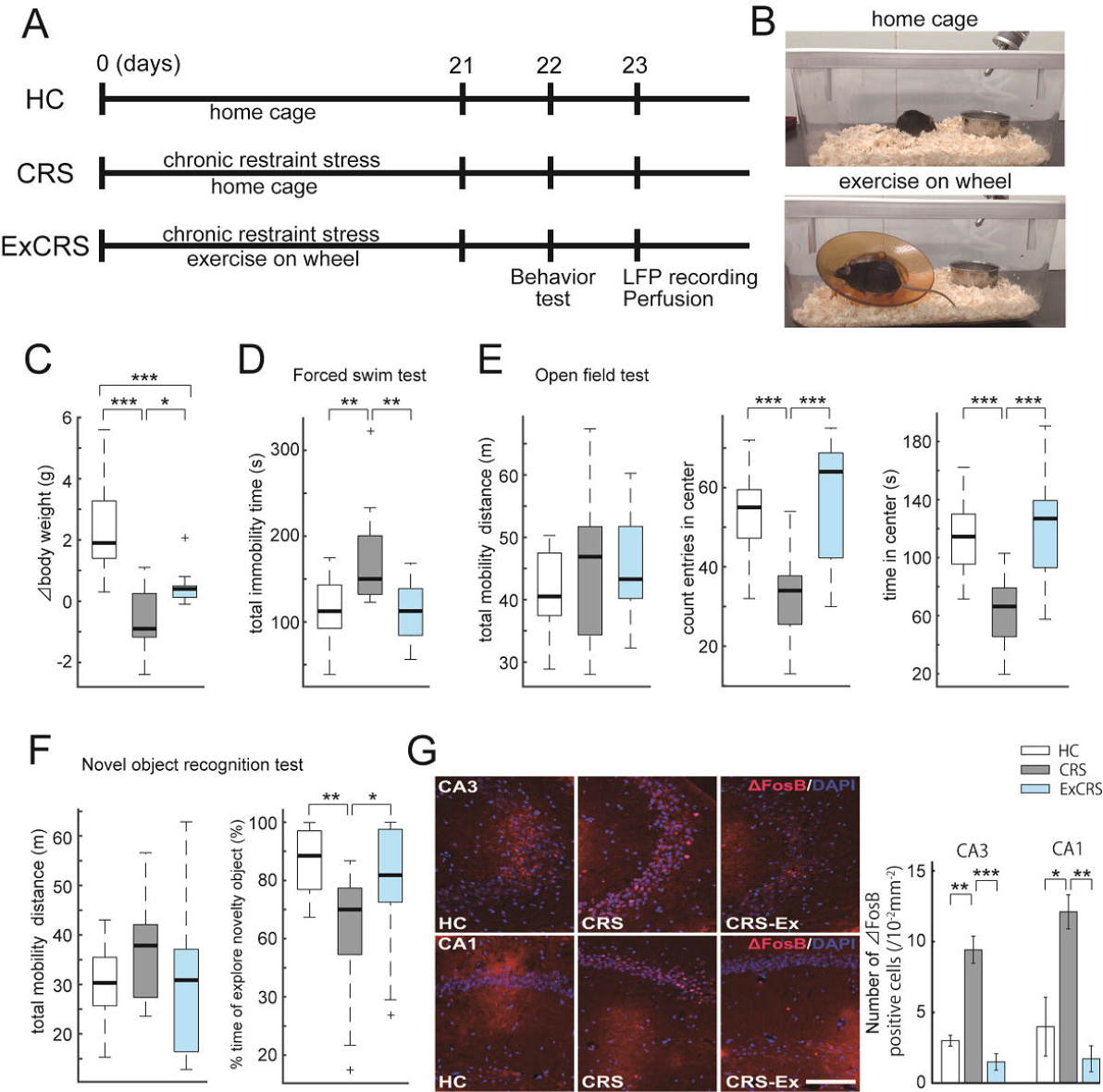
Physical exercise ameliorates depressive symptoms in the restraint stress-induced depression model of mice. **(A) Schematic of experimental procedures**. The mice were reared for 3 weeks in three conditions; 1) normal housing (HC: healthy control), 2) chronic restraint stress 4 hours a day (CRS), and 3) chronic restraint stress 4 hours a day and simultaneous physical exercise (ExCRS). **(B) The normal cage (upper) and the cage with running wheels for exercise (lower). (C) The change of body weight before and after the experiment in each group of mice. (D) Forced swim test**. The total time of immobility in 6 min was measured. **(E) Open field test in 10 min**. Left: total distance traveled. Center: Number of times the mice entered the center of the open field. Right: Total time the mice spent in the center of the open field. **(F) Novel object recognition test**. Left: Total distance the mice traveled in 10 min. Right: Percentage of time the mice spent to explore novelties in 10 min. **(G) Morphological analyses of the hippocampal activity**. Left: Representative images of immunohistological staining of ΔFosB positive cells in hippocampus CA3 and CA1 at day23. Right: Quantification of ΔFosB positive cells in CA3 and CA1. Scale bar: 100 μm. Data represent the mean ± SD. One-way ANOVA and the Tukey-HSD test (C-G). n = 15 mice / group (C-F) and n = 3 mice / group (G). * P < 0.05, ** P < 0.01, *** P < 0.001.

In addition to the stress-induced depressive behavior, the effect of restraint stress on the hippocampal neural activity and the attenuation of the effect by physical exercise were examined in the hippocampal tissues obtained from the CRS, HC and ExCRS mice. The tissue samples were immunostained with anti-ΔFosB antibody. ΔFosB, a member of the FosB family of immediate early genes, has been reported to be upregulated, responding to the environmental stress[30–32]. The number of ΔFosB expressing neurons in the CA1 and CA3 significantly increased in CRS mice compared to that in HC mice, while the number of those neurons considerably reverted in ExCRS mice (CA3; HC-CRS, p=0.001; CRS-ExCRS, p<0.001: CA1; HC-CRS, p=0.019; CRS-ExCRS, p=0.006) (Fig.1G). Taken together, the present findings demonstrate that physical exercise ameliorates the restraint stress-induced depressive behavior, cognitive decline and the hippocampal hyperactivity.

### 3.2 Hippocampal ripples are impaired in depression and retrieved by physical exercise

Local field potential (LFP) was recorded from the dorsal hippocampal CA1, using a 16-channel silicon probe under anesthesia with urethane (Fig.2A). As was previously demonstrated, the temporary ripple oscillations appeared in the stratum pyramidale in parallel with the generation of the sharp waves in the stratum radiatum[33–35]. In the present study, the effects of stress-induced depression on the occurrence and magnitude of the hippocampal ripples and their attenuation by physical exercise were examined in the depressive mice. The ripples occurred several times in 10 s during the periods of the large irregular activity in HC, CRS and ExCRS mice (Fig.2B). The frequencies of the occurrence of ripples were 22.6, 12.7, and 25.5 in 1 min in HC, CRS, and ExCRS mice, and steel-dwass analysis demonstrated significant differences (HC-CRS, p<0.05; CRS-ExCRS, p<0.05) (Fig.2C). Ripples occurred in most cases bilaterally, and in a few cases occurred unilaterally. Furthermore, the difference in frequencies of the hippocampal ripples in both sides of the brain was examined and the data showed that the frequency in either side was quite similar (Left; HC-CRS, p=0.17; CRS-ExCRS, p=0.038: Right; HC-CRS, p=0.114; CRS-ExCRS, p=0.045) (Fig.2D). Then, the duration, frequency, and amplitude of the hippocampal ripples were analyzed in the extracted ripple waveforms. The duration, which the ripples continued, was longer in CRS mice than that in HC mice, while the extension of the duration was attenuated in ExCRS mice, compared with in CRS mice (Left; HC-CRS, p=0.22; CRS-ExCRS, p=0.62: Right; HC-CRS, p=0.093; CRS-ExCRS, p=0.334) (Fig.2E). There was no significant difference in the frequency of the ripples among HC, CRS, ExCRS mice (Left; HC-CRS, p=0.745; CRS-ExCRS, p=0.737: Right; HC-CRS, p=0.358; CRS-ExCRS, p=0.942) (Fig.2F). The amplitude of the ripples in CRS mice was samaller than that in HC mice or ExCRS mice (Left; HC-CRS, p=0.244; CRS-ExCRS, p=0.521: Right; HC-CRS, p=0.213; CRS-ExCRS, p=0.69) (Fig.2G).

**Figure 2.**
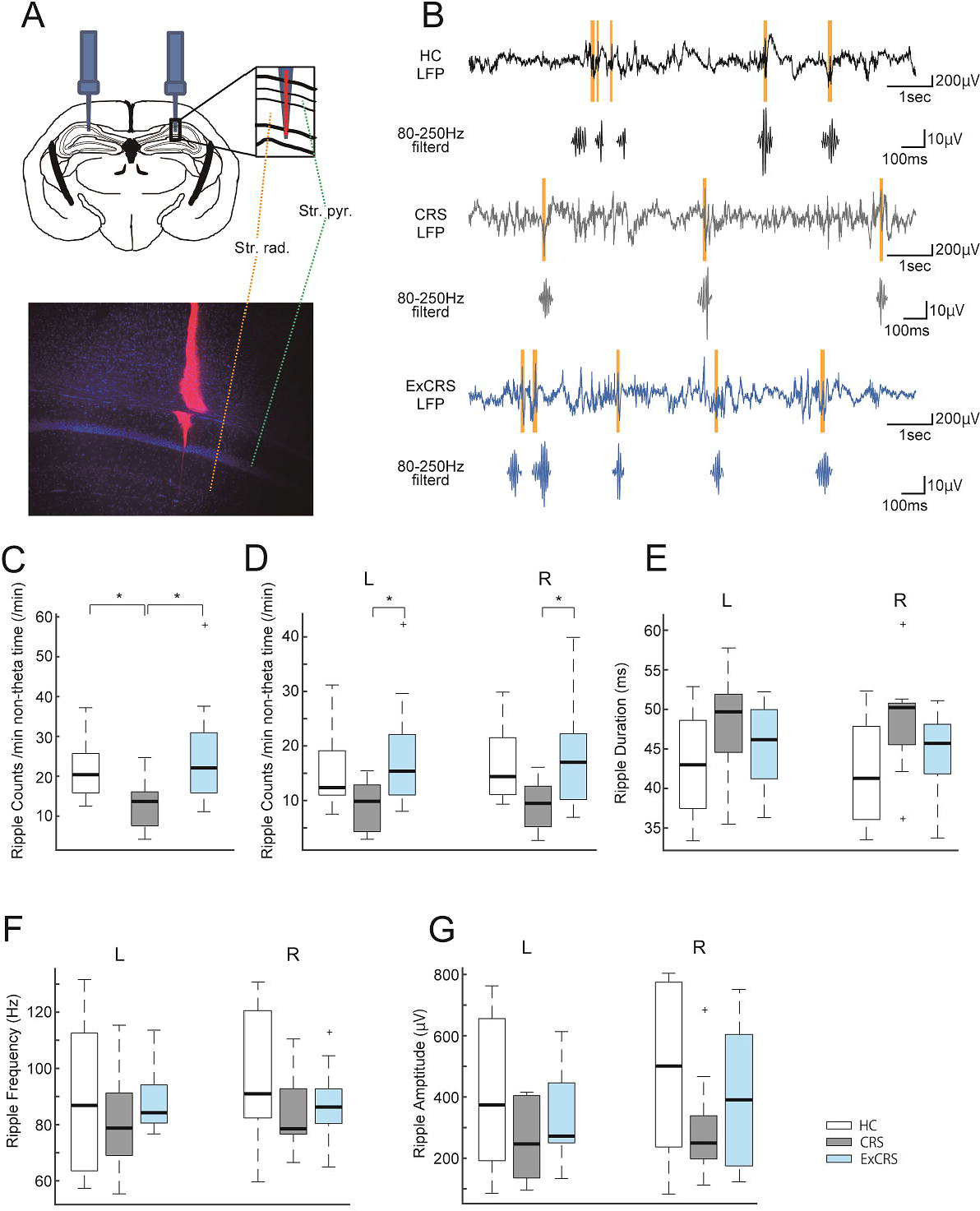
Examinations of the influence of chronic stress-induced depression on the hippocampal ripples. **(A) Schematics of LFP recording from the hippocampal CA1**. The lower panel shows an example of morphological verification of electrode location in the hippocampus. Str. Pyr.: Stratum pyramidale, Str. rad: stratum radiatum. **(B) Examples of LFP traces during non-theta periods in the CA1 stratum pyramidale**. The LFP were recorded from HC mice (top, black), CRS mice (middle, gray) and ExCRS mice (bottom, blue) in the left hemispheres. The ripples were highlighted in the LFP trace (yellow). Ripple waveforms extracted from the band-pass filtered traces were magnified and displayed below. **(C) Analysis of the number of uni-ripples. (D-G) Examinations of the influences of chronic stress on the ripples in the both hemispheres**. The numbers (D), the durations (E), the frequencies (F), and the amplitudes (G) of the ripples were examined in the left (L) and the right (R) hippocampi. One-way ANOVA and the Tukey-HSD test (D-G). n = 10 mice (HC), n = 9 mice (CRS), n = 11 mice (ExCRS) (C-G). * P < 0.05.

### 3.3 Chronic stress impairs the hippocampal theta rhythm and physical exercise restores it

As previously reported[26,36], the hippocampal LFPs alternated between theta and non-theta patterns. The non-theta period, which consists of the large irregular activity and the small irregular activity, contained the ripples (Fig.3A). The theta state was automatically detected by an algorithm based on the theta/delta power ratio of LFP in the stratum radiatum. The ratio of the theta period to the whole recorded period was 24.9, 8.2, and 38.8 (%) in HC, CRS, and ExCRS mice (n=15 respectively), and significantly reduced in depression and reverted by physical exercise (HC-CRS, p=0.045; HC-ExCRS, p=0.086; CRS-ExCRS, p<0.001) (Fig.3B). The frequency of the theta for 10 min also showed the similar tendency (HC-CRS, p=0.017; HC-ExCRS, p=0.203; CRS-ExCRS, p<0.001) (Fig.3C). The duration of the theta shortened in depression and considerably reverted by physical exercise (HC-CRS, p=0.237; HC-ExCRS, p=0.290; CRS-ExCRS, p=0.0095) (Fig.3D). Furthermore, the gamma oscillations, contained in the theta LFPs in the stratum radiatum, were examined. The powers of the slow gamma (40- 55 Hz) and fast gamma (65- 90 Hz) oscillations were calculated by spectral analysis (Fig.3E). The spectral powers of the theta, slow gamma, and fast gamma among HC, CRS, and ExCRS mice were compared. The theta power was significantly lower in CRS mice than that in HC mice, while reverted in ExCRS mice (Left; HC-CRS, p=0.102; CRS-ExCRS, p=0.94: Right; HC-CRS, p=0.234; CRS-ExCRS, p=0.074, Fig.3F). Interestingly, the theta power reverted in the right hippocampus dominantly. On the other hand, the slow and fast gamma powers were significantly lower in CRS mice than those in HC mice (fast gamma Left, p=0.040; fast gamma Right, p=0.049; slow gamma Left, p=0.035; slow gamma Right, p=0.022) (Fig.3G). Of note is that in the right hippocampus the fast gamma dominantly reverted as the result of exercise in ExCRS, while that reverted less in the left hippocampus (fast gamma Left, p=0.888; fast gamma Right, p=0.111; slow gamma Left, p=0.544; slow gamma Right, p=0.027).

**Figure 3.**
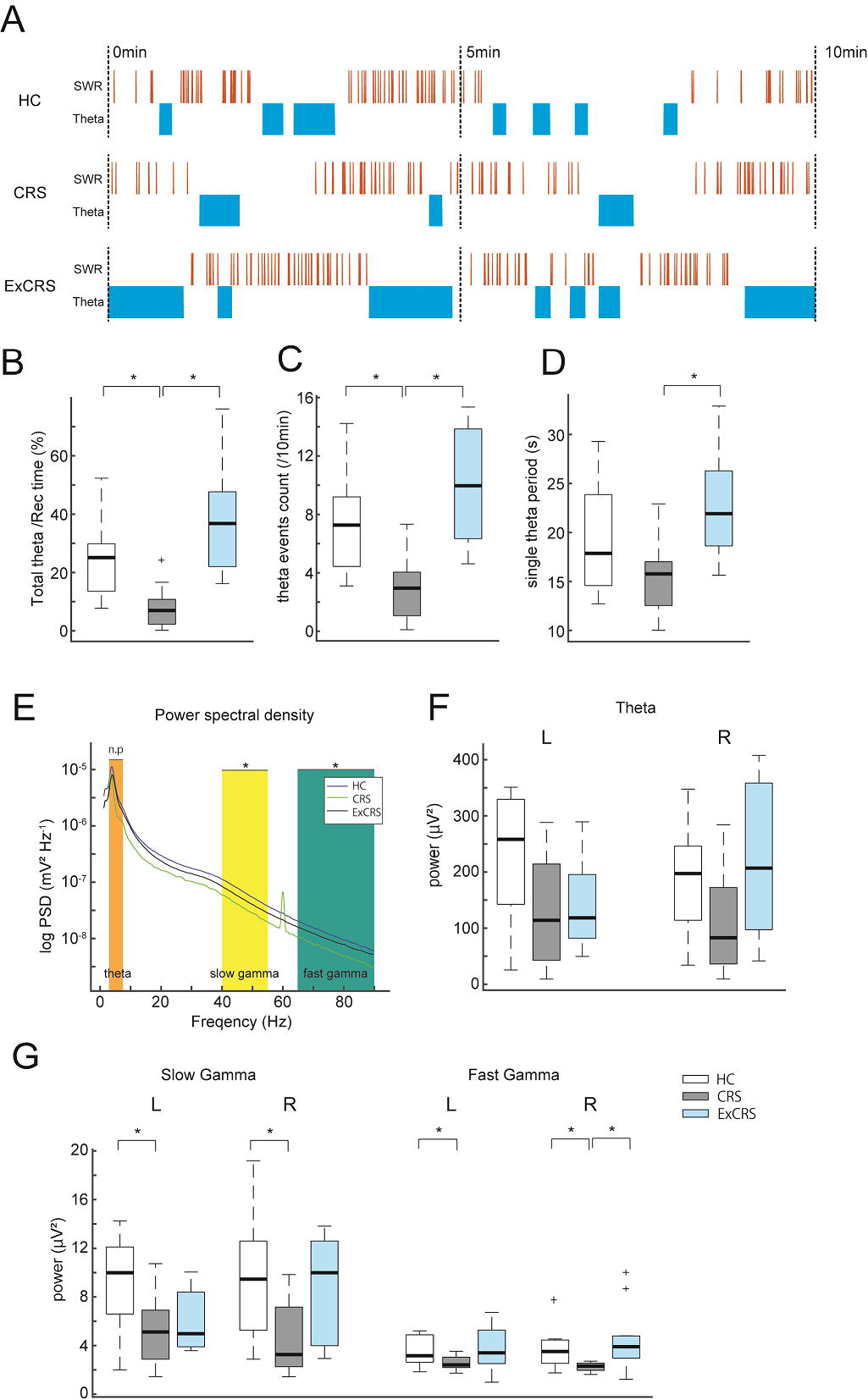
Examinations of the defects of the theta and the theta-associated gamma oscillations in the CA1 stratum pyramidale in the depression model of mice. **(A) Schemes of the typical sharp wave ripples (SWR) and theta rhythms**. The SWRs are shown in orange, and the theta rhythms are shown in blue. **(B-D) Analyses of the influence of chronic stress on the theta**. Statistics for spontaneously occurring theta periods are computed for (B) the proportion of theta periods in the total recording time, (C) the frequency of theta state, (D) the duration of single theta period. **(E) Power spectral densities of LFP in the CA1 stratum radiatum**. Shaded areas represent the frequency ranges for slow (40–55 Hz) and fast (65–90 Hz) gamma oscillations. **(F, G) Analyses of the influence of chronic stress on the theta and gamma powers**. (F) The theta power in HC, CRS, and ExCRS mice. (G) The slow and fast gamma power in the left (L) and right (R) hemispheres. One-way ANOVA and the Tukey-HSD test (B-G). n = 9 mice (CRS), n = 11 mice (ExCRS) (B-G). *P < 0.05.

### 3.4 Physical exercise attenuates the defect of the hippocampal adult neurogenesis

As was demonstrated in previous studies, the defect of hippocampal adult neurogenesis is involved in the etiology of depression[37–39]. In the present study the influence of physical exercise on the hippocampal adult neurogenesis was examined in the depression model of mouse. The hippocampal tissues were obtained from the mice and their neural stem cells were immunostained with anti-Sox2 antibody. In CRS mice, the number of neural stem cells in the DG was considerably fewer than that in HC mice, while in ExCRS mice the number of neural stem cells restored (HC-CRS, p=0.205; CRS-ExCRS, p=0.022) (Fig. 4A and B).

**Figure 4.**
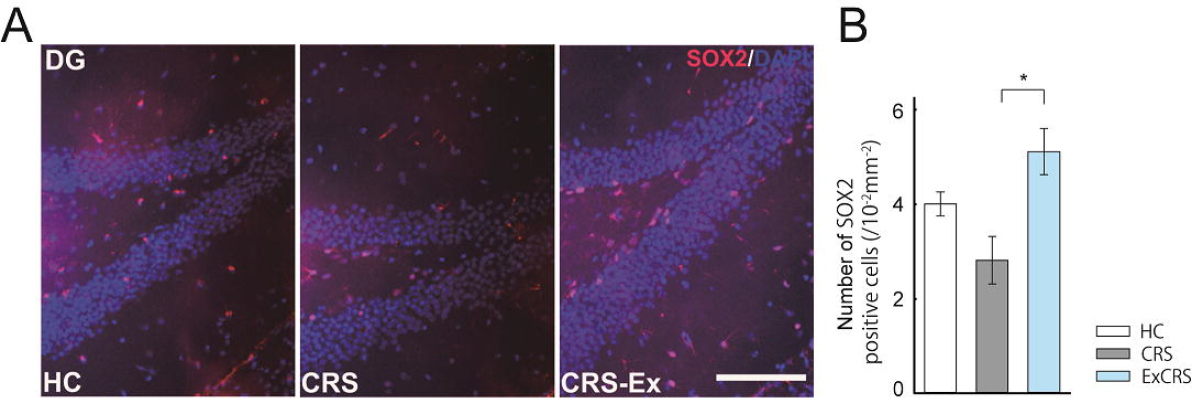
Physical exercise ameliorates the defect of hippocampal adult neurogenesis in depression. **(A) Representative images of Sox2-expressing neural stem cells in the hippocampal dentate gyrus (DG) at day 23**. Scale bar: 100 μm. **(B) The defect of adult neurogenesis in depression and its amelioration by physical exercise**. Data represent the mean ± SD. * P < 0.05.

### 3.5 Physical exercise suppresses stress-induced neuroinflammation in the hippocampus

Accumulated evidence demonstrated that the inflammation with microglial activation in the hippocampus is correlated with the dysfunction of hippocampal adult neurogenesis and implicated in the etiology of depression[40,41]. In the present study microglial activation, which accompanies the morphological change from ramified to amoeboid shape and the change in length and number of processes[28,29], was examined in the brain tissue samples of the depression model mouse. The change of microglial shape was morphologically analyzed by immunohistochemical staining with anti-Iba1 antibody, and the number and total length of microglial processes were quantified by skeleton analysis of ImageJ plug-in. There was no significant difference between HC, CRS and ExCRS mice in the number of microglia in the hippocampal DG, CA1 and CA3 (Fig. 5A and C), while the number and total length of microglial processes in CRS mice were significantly less than those in HC mice in the DG, CA3 and CA1 (number of processes: DG, p=0.005; CA3, p=0.024; CA1, p=0.009. length: DG, p=0.019; CA3, p=0.019; CA1, p=0.026). Further analyses revealed that the number and total length of microglial processes significantly increased in ExCRS mice, whose microglia took ramified shape, compared to those in CRS mice (number of processes; DG, p=0.001; CA3, p=0.014; CA1, p=0.006: total length; DG, p<0.001; CA3, p=0.004; CA1, p=0.003) (Fig.5B, D and E).

**Figure 5.**
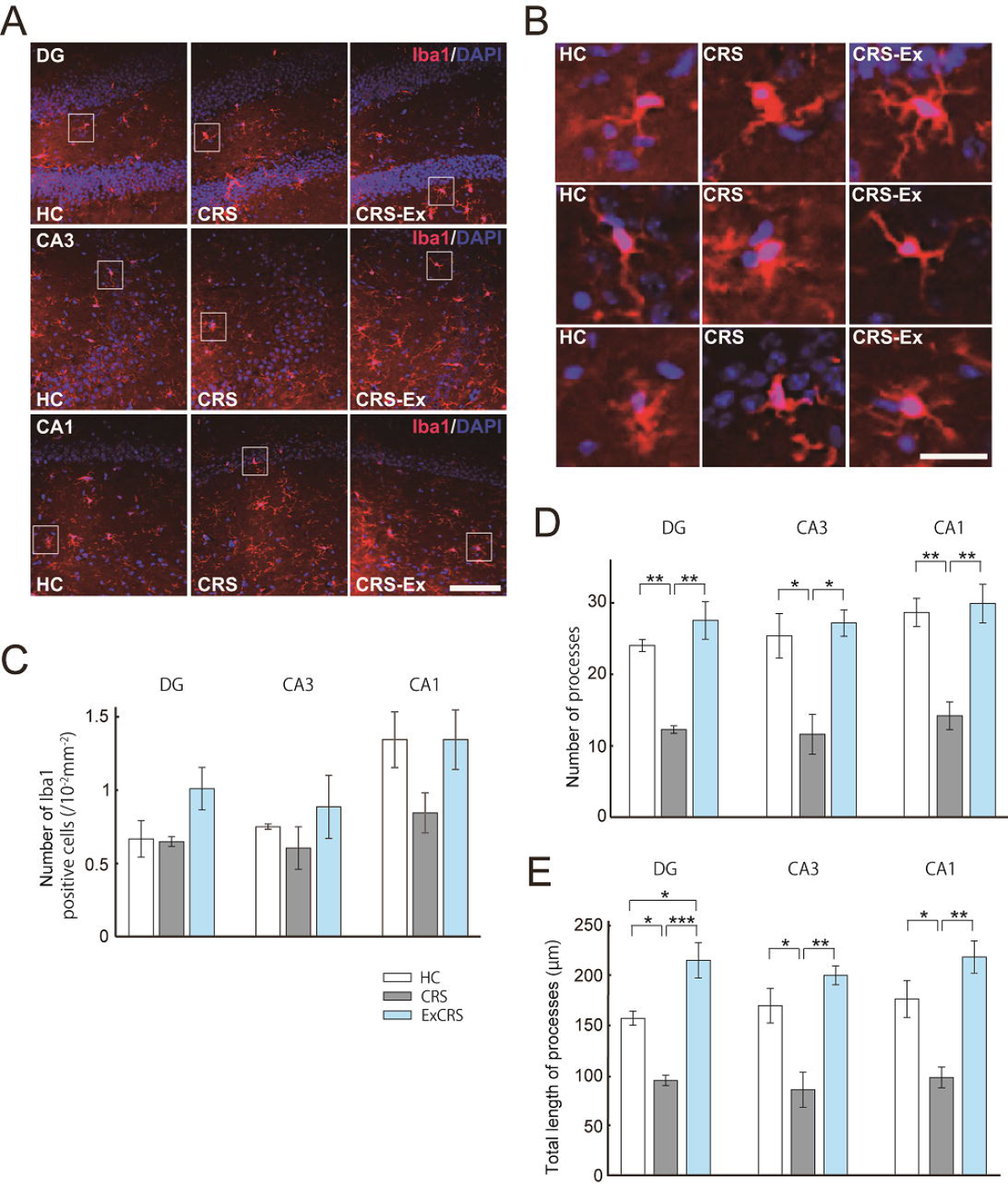
The suppression of neuroinflammation in depression by physical exercise. **(A) Representative images of Iba1-expressing microglia in the hippocampal DG, CA3 and CA1 in mice at day 23**. Scale bar: 100 μm, **(B) Magnified images of the boxed areas in (A)**. Scale bar: 50 μm. **(C-E) Morphological analyses of the hippocampal microglia**. (C) The number of microglia, (D) the number of processes in a cell, (E) the total length of processes in a cell. DG; dentate gyrus. Data represent the mean ± SD. One-way ANOVA and the Tukey-HSD test (C-E). n = 3 mice / group (C-E). * P < 0.05, ** P < 0.01, *** P < 0.001.

## 4. Discussion

The beneficial effect of physical exercise on the amelioration of depressive symptom has been well appreciated, but neural bases underpinning its effect remains to be elucidated. In addition, the neurophysiological indexes to estimate the severity of depressive symptom in the live brain have not been identified. Recent findings revealed that the dysfunction of the hippocampal neural connections with the cerebral cortex, especially prefrontal cortex, amygdala and hypothalamus are involved in the etiology of depression, and in addition, hippocampal SWRs was reported to correlate with the disease condition of depression. Therefore, in the present study the alteration of hippocampal SWRs in association with depressive symptom was examined by developing the restraint stress-induced mouse model of depression.

In mouse model of depression, physical exercise ameliorated depressive symptoms such like body weight loss, reduction of motivation, anxiety, cognitive impairment (Fig. 1A-F). In addition to those psychiatric defects, hippocampal malfunction with neural hyperactivity, correlated to the neuropathology of depressive behavior, was detected in the model mice (Fig. 1G). Those findings demonstrate that physical exercise diminishes the influence of restraint stress and relieves the depressive symptoms.

The present data suggested that hippocampal SWRs, which were previously demonstrated to correlate with cognitive and emotional processing in the cortico-hippocampal connections[42], were affected in the pathological condition of depression. Furthermore, physical exercise diminished the effect of restraint stress on numbers, duration, and frequency of ripples (Fig. 2C-G). Despite our previous observation, in which the hippocampus on each side of the brain was demonstrated to asymmetrically process cortical inputs[43], hippocampal SWRs in either side of the brain didn’t generate differential pattens (Fig. 2C-G).

The influence of restraint stress-induced depression on the hippocampal theta rhythm, which alternates with hippocampal SWRs and underlies several aspects of cognition such as attention and exploration, and the therapeutic effect of physical exercise on it were examined. The findings demonstrated that exercise restored the theta’s disruption and the impairment of gamma rhythm, which has been previously reported to correlate with working memory (Fig. 3B-G). The hippocampal theta rhythm reflects the input from the hypothalamus and its dysfunction results in the defects of motivation[44]. Its restoration, therefore, leads to amelioration of depressive symptom.

Accordingly, the present findings regarding the effects of exercise on the restoration of the hippocampal electrophysiological activity and their correlation with the attenuation of depressive symptoms, imply that electrophysiological indexes of hippocampal activity such as SWRs or theta rhythm can be utilized for the neuropathological markers of depression.

As previous studies showed, dysfunction of the hippocampal adult neurogenesis and causative inflammation with microglial activation occurred in our restraint stress induced-depression model, in which physical exercise attenuated those pathological conditions (Fig. 4 A, B and Fig. 5 A-E). Inflammation and consequent defect of the hippocampal adult neurogenesis are involved in the environment-responding reconstruction of neural circuit and might affect the hippocampal activity expressed as SWRs and theta rhythm. Because dysfunction of the hippocampal adult neurogenesis is difficult to be non-invasively accessed, alternative approach such like estimation of SWRs or theta rhythm is more practical for diagnosis of depression.

In conclusion, the present study demonstrates the correlation of electrophysiological indexes, hippocampal and theta rhythm, with the pathophysiology of stress-induced depression, and the possible application of those indexes for diagnosis of depression. The current data shows that the dysfunction of hippocampal connections with several areas of the brain functioning in cognition, emotion, motivation, such as the cerebral cortex, amygdala, hypothalamus, underlies the etiology of depression, and therefore, the development of diagnostic and therapeutic methods, targeting the hippocampal circuits might open the new avenue for the cure of depression.

